# Afrotropical Tree Communities May Have Distinct Responses to Forecasted Climate Change

**DOI:** 10.1101/823724

**Authors:** Chase L. Nuñez, James S. Clark, John R. Poulsen

## Abstract

More refined knowledge of how tropical forests respond to changes in the abiotic environment and human disturbance is necessary to mitigating climate change, maintain biodiversity, and preserve ecosystem services. To evaluate the unique response of Afrotropical forests to changes in the abiotic environment and disturbance, we employ species inventories, remotely sensed historic climatic data, and future climate predictions collected from 104 1-ha plots in the central African country of Gabon. We forecast a 3 - 8% decrease in Afrotropical forest species richness by the end of the century, in contrast to the 30-50% loss of plant diversity predicted to occur with equivalent warming in the Neotropics. This work reveals that community forecasts are not generalizable across regions, and more representative studies are needed in understudied biomes like the Afrotropics. This study serves as an important counterpoint to work done in the Neotropics by providing contrasting predictions for Afrotropical forests with substantially different ecological, evolutionary, and anthropogenic histories.

## Introduction

The anticipated pace of global warming is predicted to result in large declines of tropical biodiversity, leading to biotic attrition of the lowland tropics (Sala *et al.* 2000; van Vuuren *et al.* 2006; Colwell *et al.* 2008; Feeley *et al.* 2011; Hooper *et al.* 2012; Dexter *et al.* 2018, but see Feeley and Silman 2010). Effects of climate change may be most acutely felt by long-lived organisms like trees that endure several degrees of warming within a single lifetime, without the benefit of adaptation available to shorter lived species (Malhi *et al.* 2014). Tropical forests contain over 40,000 tree species (Slik *et al.* 2015), shelter over half of all animal species (Pimm and Raven 2000), and store much of the planet’s carbon (Sullivan *et al.* 2017) while covering only 7% of the Earth’s surface (Corlett and Primack 2011). Despite their outsized value, tropical forests are notably understudied, and most *in situ* species inventory data come from a few intensively studied sites in the Neotropics (Schimel et al. Global Change Biology 2015). Neotropical studies demonstrate that community composition and function are degrading in response to climate change (Engelbrecht *et al.* 2007; Bongers *et al.* 2009; Poorter *et al.* 2017), with early successional species thriving in warmer soil temperatures at the expense of late-successional species that require cooler microhabitats (Colwell *et al.* 2008). These processes have contributed to predictions of 30-50% loss of plant diversity with a 5°C temperature increase for most South American tropical forests (Colwell *et al.* 2008; Feeley and Silman 2010).

It is unclear whether the world’s other tropical regions will respond similarly to climate change (Malhi and Wright 2004; Maslin *et al.* 2005; Parmentier *et al.* 2007; Malhi *et al.* 2013; Mayaux *et al.* 2013; Enquist *et al.* 2017; Sullivan *et al.* 2017). Afrotropical forests are distinct in having comparatively few wet-affiliated species given their climate (Leal 2009), and a high proportion of large trees that grow and recolonize rapidly (Fayolle *et al.* 2012; Gond *et al.* 2013). These differences in community-level traits are hypothesized to have arisen from Africa’s unique climatic past (Haffer 1969; Maley 1996; Maslin *et al.* 2005; Oslisly *et al.* 2013; Willis *et al.* 2013). In direct contrast to the Neotropics, abnormally cool and dry conditions during the last glacial maximum reduced Afrotropical forests to small fragmented patches (Walker and Sleath 2017). This may have selected for species able to survive extreme aberrations in temperature and precipitation and then quickly disperse from refugia to recolonize the landscape cleared by receding glaciers (Leal 2009), potentially making them more resilient to climate change than their Amazonian or Asian counterparts (Hansen and DeFries 2004; Gardner *et al.* 2007). Indeed, a pair of studies comparing changes in canopy structure found few lingering effects of drought on Afrotropical forest canopy (Asefi-Najafabady and Saatchi 2013), but did find lingering canopy effects of drought on southwest Amazonia (Saatchi *et al.* 2013). Although it is clear that Neotropical species responses to climate change are unlikely to be an adequate proxy for Afrotropical forests, no landscape-scale predictions of Afrotropical community responses to climate have been made. This work is urgently needed—the current climate of Equatorial Africa is already near the lower temperature-precipitation threshold of rainforest viability (Malhi and Wright 2004; Pan *et al.* 2011), after which a rapid shift in the ecosystem could occur (Willis *et al.* 2013).

To evaluate how Afrotropical tree species will respond to future climate change, we model tropical forest tree species distributions using a large array of tree plots, remotely sensed historic climatic data, and future climate predictions for the densely forested central African country of Gabon. This first-of-its-kind study serves as an important counterpoint to work done in the Neotropics by providing contrasting novel predictions for Afrotropical forests with substantially different ecological, evolutionary, and anthropogenic histories. We hypothesize that the disturbance-rich past of Afrotropical communities will result in two divergent responses to forecasted climate change: 1) Afrotropical community forecasts will demonstrate lower levels of species loss than the 30-50% loss predicted for the Neotropics; and 2) early-successional species will increase in number at the expense of declining abundance of late-successional species.

**Figure 1:**
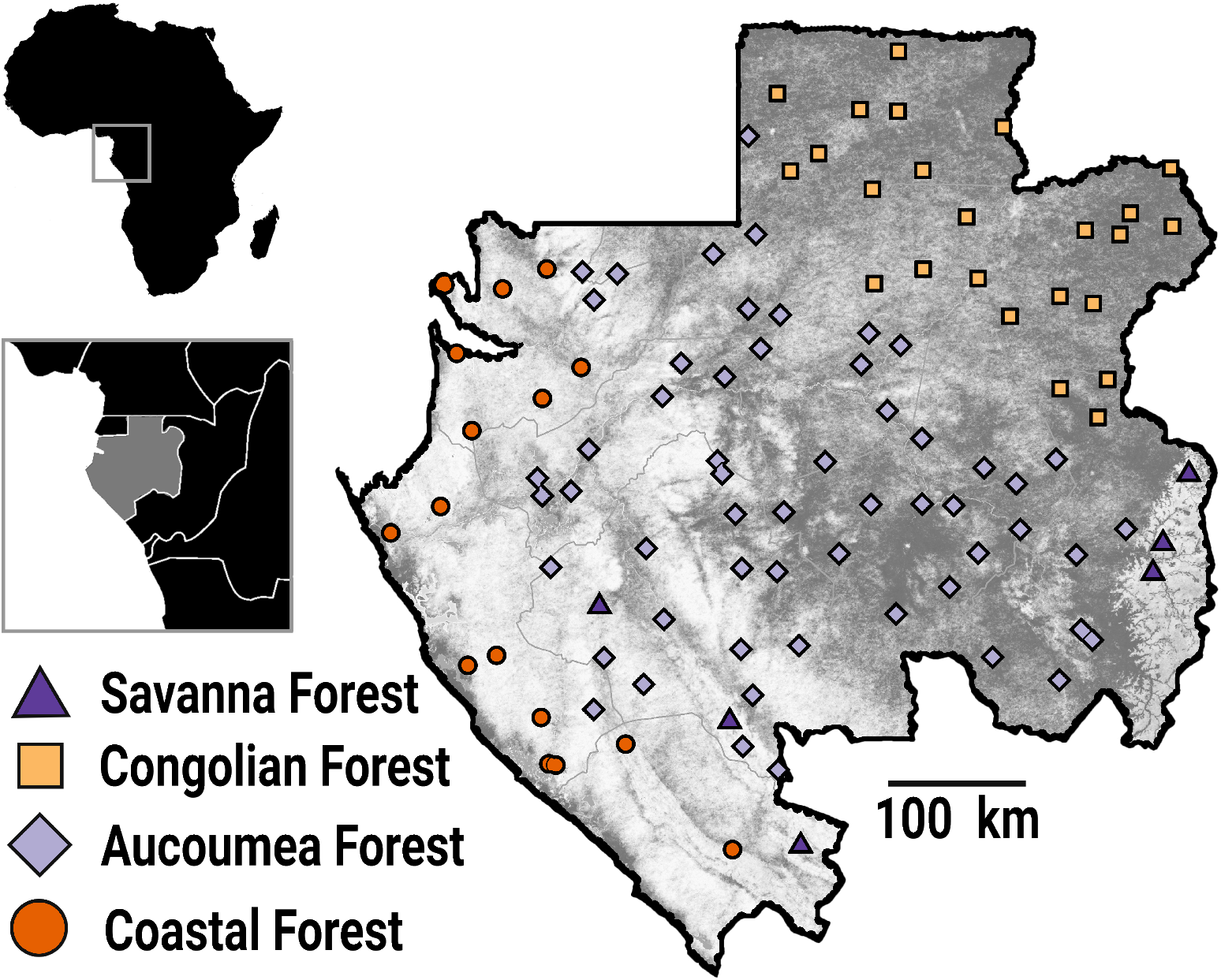
Map of all 104, 1-ha inventory plots located in the Central African country of Gabon. Plots were located by a systematic-random design to capture the full breadth of Gabon’s forest types and environmental conditions. Shapes indicate one of four possible forest-types of each plot: Savanna Forest (triangles), Congolian Forest (squares), Aucoumea Forest (diamonds), or Coastal Forest (circles).

## Methods

### Tree Inventory Data

In this study, we employ tree census data from Gabon’s National Resource Inventory—a national network of tree plots for estimating forest biomass and carbon stocks (Figure 13) (Carlson *et al.* 2017; Poulsen *et al.* 2017; Wade *et al.* 2019). Gabon is the second most forested country in the world, with a forest cover of 87%, a deforestation rate near zero (Sannier *et al.* 2014), and one of the highest densities of carbon in Central Africa (Saatchi *et al.* 2011). Between 2012 and 2013, trained technicians established 104 1-ha forest plots based on a stratified random sampling design that consisted of dividing the country into 100 50 × 50 km cells and randomly locating a sample site within each of the cells. This design ensured an unbiased sampling of Gabon’s forest (40.4% old-growth, 28.8% logged, 30.8% secondary) and edaphic types (69.2% terra firma, 22.1% seasonally flooded forest, 8.6% swamp). Every tree with a diameter-at-breast height (DBH) ≥ 10 cm was mapped, measured and identified to species by trained field teams following standard protocols for plot establishment and measurement (Phillips and Baker 2002). 621 species from 296 genera were catalogued. We analyzed data for all species that occurred on at least 30 of the 104 plots, resulting in 34,460 stems representing 76 of the most widely occurring species. The five most common taxa (31% of all stems) were *Santiria trimera, Dichostemma glaucescens, Plagiostyles Africana, Aucoumea klaineana*, and *Diospyros spp.* The five least common taxa included in the model (1% of all stems) were *Duboscia macrocarpa*, *Ongokea gore*, *Zanthoxyllum heitzii*, *Klainedoxa sp.*, and *Erythrophleum ivorense*. Each species was assigned, if possible, into an “early-successional” or “late-successional” class based on growth form, available trait data, and habitat class (Whitmore 1989; Raich and Khoon 1990; Finegan 1996; Davies and Semui 2006; Chazdon *et al.* 2010; Hardwick and Elliott 2016).

### Climate Data

Long term average historical precipitation and temperature data for each plot were derived from the NASA TerraClimate product (Abatzoglou *et al.* 2018) for 1985 - 2017, accessed using Google Earth Engine (Gorelick *et al.* 2017). We derived projected precipitation and temperature data for each plot using the NASA Earth Exchange Global Daily Downscaled Projections (NEX-GDDP) database at a resolution of 0.25 degrees (~25 km × 25 km). This dataset provides downscaled projections for two of the most commonly used Representative Concentration Pathways (RCP 4.5 and RCP 8.5) from the 21 General Circulation Models that were produced and distributed under the Coupled Model Intercomparison Project Phase 5 (CMIP5). All models of temperature agree that Gabon will continue to warm through the end of the century; thus, within each RCP scenario, we took the ensemble mean prediction of all 21 CMIP5 GCMs to forecast of forest response (Figures 14A, 14B). Models of precipitation disagree as to whether precipitation will increase or decrease, producing an ensemble mean that shows no substantive change by 2099 (Figure 14C). This is due in large part to a lack of rain gauges in Central Africa (Washington *et al.* 2013), and also from the northward shift of the inter-tropical convergence zone resulting from ocean-driven atmospheric circulation shifts (James *et al.* 2013). To acknowledge the uncertainty in whether the dry or wet models are more plausible, here we forecast community change given both a wet and a dry scenario. The wet scenario takes the mean prediction of the five models predicting the greatest increase in precipitation, while the dry scenario uses the mean prediction from the five models predicting the greatest decreases in precipitation (Appendix A3). Although representing the extreme scenarios, this forecasted climate space is well represented by historical climate space (Appendix A6).

**Figure 2:**
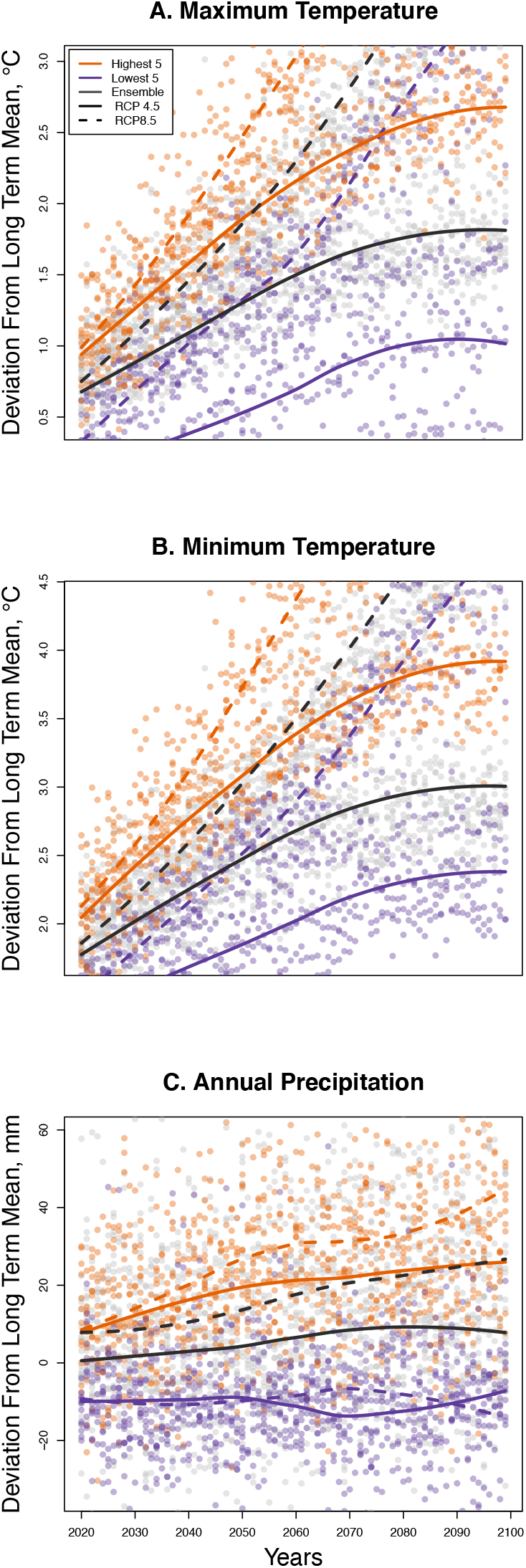
A comparison of average predicted maximum temperature (A), minimum temperature (B), and precipitation (C) in Gabon from the highest five models (orange), lowest five models (purple), and all model ensembles (grey) for RCP 4.5 (solid lines) and RCP 8.5 (dashed lines).

### Community Composition Analysis

We use a generative Generalized Joint Attribute Model (GJAM) that predicts species abundance at the scale and context used to fit the model jointly, *i.e.* on the community scale (Clark *et al.* 2017). GJAM estimates can therefore be interpreted on the scale of the observations, accounting for sample effort. Full model specifications are available from Clark *et al.* (2017). Parameters in the model include matrices of coefficients B relating X to Y and the residual species covariance matrix Σ. In effect, Σ represents the covariance between species beyond what is explained by environmental covariates. This variation can come from interactions between species, unmeasured environmental variables, and other sources of error. The likelihood is: [Y_1_,…,Y_S_ |X, B, Σ], where subscripts refer to species 1 through S. Model fitting is done on the observation scale, and is based on the posterior distribution, [B, Σ |X,Y]∝[Y1,…,YS |B, Σ][B, Σ]. The right-hand side of the equation is the likelihood and the prior distribution, [B, Σ], which is non-informative.

### Forecasting Species Composition

The predictive distributions combine the posterior parameter estimates calibrated from long-term climate data with a prediction grid of forecasted covariate values (precipitation and temperature). *X\** is a vector of environmental covariates that generate a response vector of species *Y*\*= *Y*_1_, …, *Y*_*S*_, and (*X*\*, *Y*\*) is a pair of vector observations used to fit the model. The predictive distribution [*Y*\*|*X*\*] = ∫ [*Y*\* | B, Σ, *X*\*][B, Σ |*X*, *Y*]*d*(B, Σ) is obtained by Monte Carlo integration. The two factors in the integrand are the likelihood and posterior distribution. To quantify how changes in climate will affect total species counts regardless of forest type, we first run the model using forest type (Congolian, savanna, coastal, Acoumea) as a random effect. We then use forest type as a factor to make predictions about species counts within forest types. Covariates used to fit the model were limited to those for which there are predictions available from the NASA NEX GDDP GCM’s for years 2020-2099 and having variance inflation factors less than 3. Model estimates were taken from 100,000 iterations, discarding the first 1000 iteration as pre-convergence. We visually inspected trace plots to confirm convergence and adequate mixing (Appendix A5) and validated model fit by comparing predicted and observed discrete species abundances (Appendix A7).

## Results

Of the 76 most abundant species analyzed, species richness in Gabon’s forests is projected to decrease by 3 - 8% by the end of the century, although there is variation among precipitation models and RCP scenarios (Figures 15, 16, Appendix A1). The dry model predicts a loss of 2.58 focal species per plot ± SE 5.36 in the low emissions scenario (RCP 4.5, Figure 16A), and −5.85 ± 5.12 species in the high emissions scenario (RCP 8.5, Figure 16C). The wet model predicts fewer species losses (RCP 4.5: −3.17, ± 5.37; RCP 8.5: −5.96 ± 5.11, Figures 15A, 15C). Species losses were consistent across forest types (Figures 15B/15D, 16B/16D, Appendix A2) with greatest losses in the RCP 8.5 scenario (Figures 15D, 16D) compared to RCP 4.5 (Figures 15B, 16B). Of the 76 species we analyzed approximately one third are predicted to increase in abundance and two-thirds of species likely to decrease in abundance. The most likely species to increase in abundance include *Diospyros spp*., *Aucoumea klaineana*, and *Staudia gabonensis*. By contrast, *Santira trimera*, *Plagiostyles africana*, and *Dichostemma glaucescens* are predicted to decrease in abundance (Figure 17). Net change in predicted species abundance was similar between wet and dry models and among RCP scenarios (Figure 17, Appendix A4).

**Figure 3:**
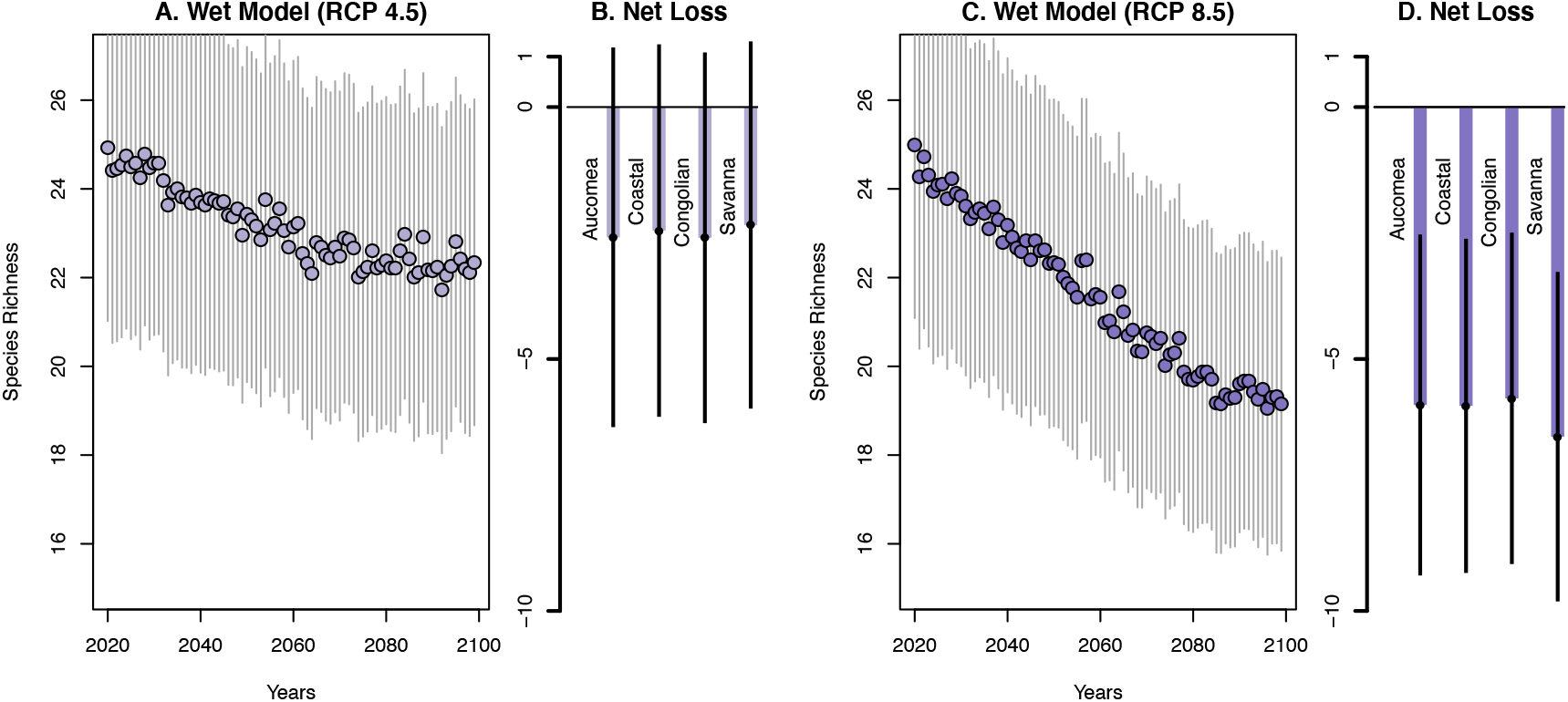
Declines in estimated plot-level species richness and the estimated number of lost species for each forest type by the end of the century for RCP 4.5 (A, B) and RCP 8.5 (C, D) for the wet model.

**Figure 4:**
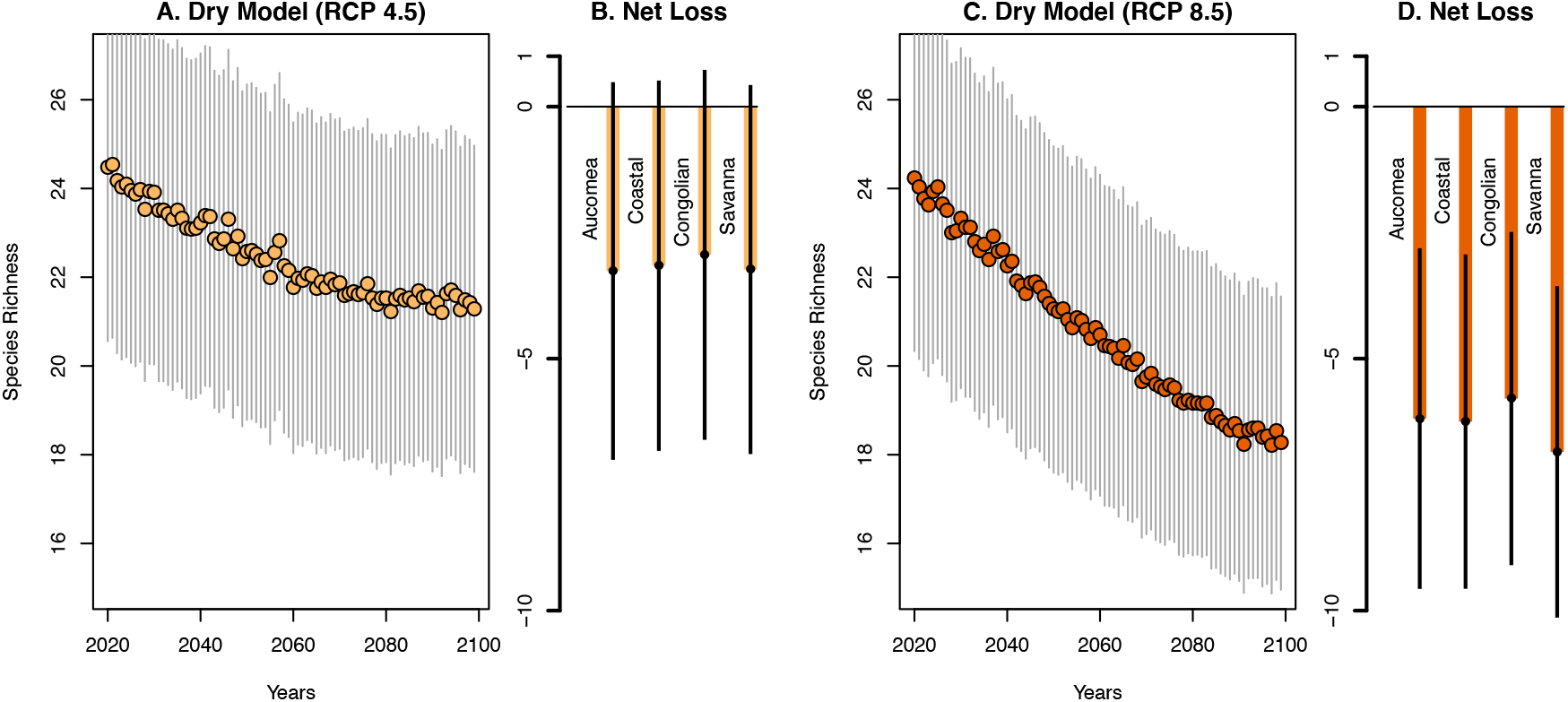
Declines in estimated plot-level species richness and the estimated number of lost species for each forest type by the end of the century for RCP 4.5 (A, B) and RCP 8.5 (C, D) for the Dry model.

**Figure 5:**
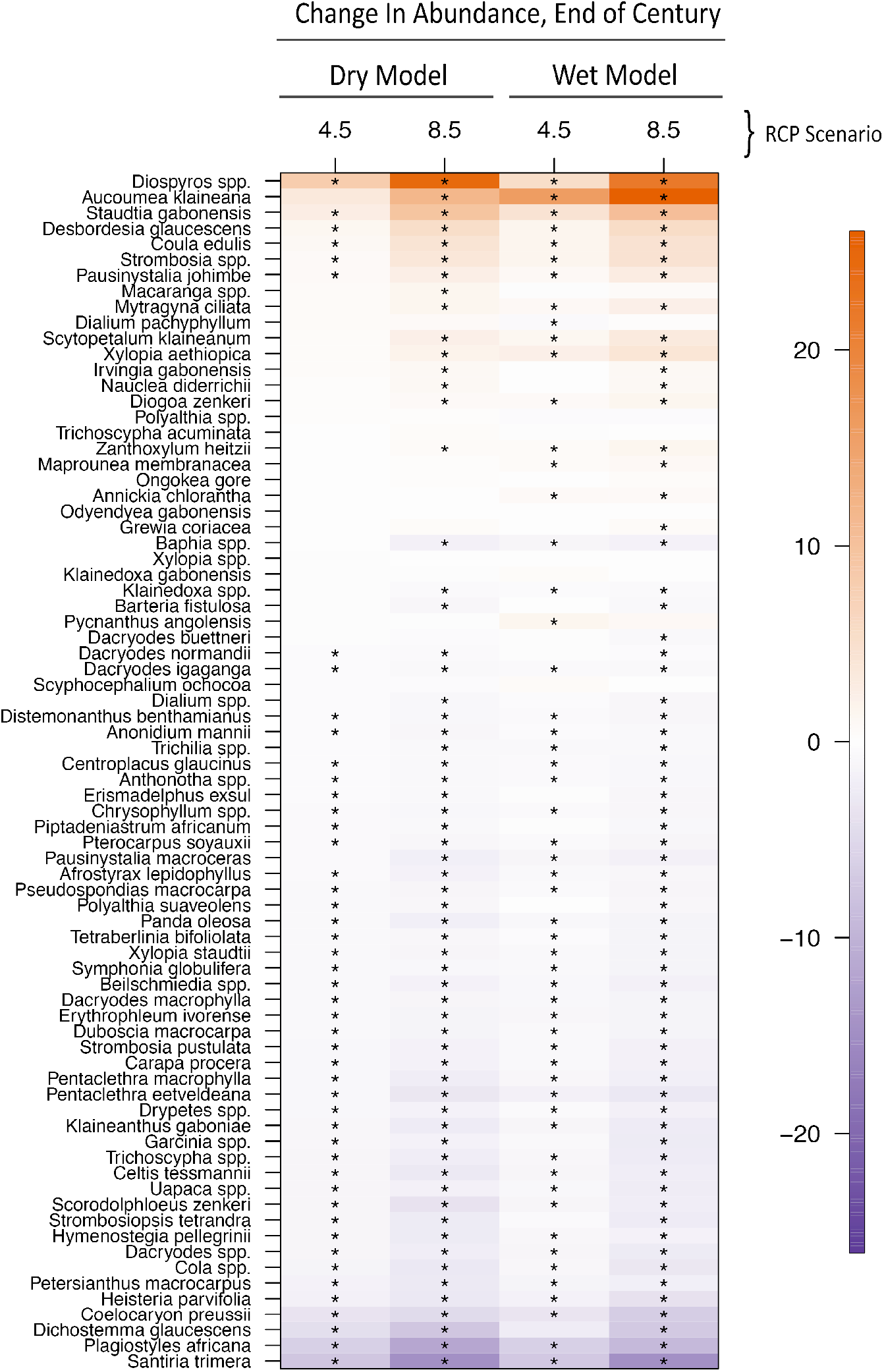
Predicted change in abundance for each species (y axis) by the end of the century for RCP 4.5 and RCP 8.5 in both the ‘wet’ and ‘dry’ model ensembles (x axis). Box colors correspond to estimated changes in species abundance. “*” symbol in box denotes species-model predictions that do not overlap with 0.

## Discussion

We demonstrate a 3 – 8% decrease in Afrotropical forest species richness by the end of the century for the most abundant tree species. This prediction is substantially less severe than 30-50% predicted for the Neotropics (Colwell et al. 2008; Feeley and Silman 2010), and lends support to the argument that the unique evolutionary past of Afrotropical forest communities (Malhi and Wright 2004; Maslin *et al.* 2005; Parmentier *et al.* 2007; Malhi *et al.* 2013; Mayaux *et al.* 2013; Enquist *et al.* 2017; Sullivan *et al.* 2017) could have made them more resilient to climate change than their Amazonian or Asian counterparts (Hansen and DeFries 2004; Gardner *et al.* 2007). It also indicates that predictions of tree species responses to climate change are not generalizable across continents. Surprisingly, we find that losses do not differ appreciably among forest types, indicating that no habitats will be immune from species changes. The equivalent responses may also be a result of the subjective distinction between our four forest types – communities are complex assemblages of species that may not always readily lend themselves to clear distinctions.

We find that species-specific changes in abundance varied considerably, with approximately a third of all species likely increasing in abundance and two-thirds of species likely decreasing in abundance. Our hypothesis that disturbance-related species would increase at the expense of climax species was not well supported. Of the ten species most likely to *increase in abundance*, only two are considered early successional. Of the ten species most likely to *decrease in abundance*, five were early successional. It is interesting that the species increasing are not just those that are fast growing, low wood density species associated with disturbance. For example, two of the species most likely to increase are *Acoumea klaineana* (a light-loving, low wood density species), and *Diospyros spp*, the genus that incudes ebony (high wood density, slow growing). Among the species expected to decrease in abundance is *Santira trimera*, one of the most widespread species throughout West and Central African rainforests, often in moist secondary forests or along rivers. Also predicted to decrease is *Dichostemma glaucescens*, a small slender tree prone to climbing other trees. These species-specific responses suggest that rather than functional groups responding similarly, tropical species respond individualistically to tropical climate change. This pattern agrees with past research (Bush 2002; Bush *et al.* 2004) and may arise from unique relationships to unmeasured abiotic variables that contribute to its response to disturbance (Nuñez *et al.* 2019). A comparison of the forecasts produced by both dry and wet models yields surprising little difference in predicted species richness. Although the total water available to trees are the product of both precipitation and temperature, these results suggest that species will respond more strongly to increases in temperature, not precipitation as it is in the Neotropics. For this reason, the increasing radiative forcing associated with RCP 8.5 resulted in greater species loss than the more conservative RCP 4.5 scenario. Temperature specific responses are consistent with the theory that a unique ecological history in the Afrotropics (Malhi and Wright 2004; Maslin *et al.* 2005; Parmentier *et al.* 2007; Malhi *et al.* 2013; Mayaux *et al.* 2013; Enquist *et al.* 2017; Sullivan *et al.* 2017) cultivated tree communities with few wet-affiliated species(Leal 2009)

Several considerations are necessary to situate this study in the literature. First and foremost, this analysis considered only the 76 most common species across plots. Rare species (occurring on fewer than 30/104 plots) have few observations and provide insufficient information on their relationship with environmental predictors to make accurate predictions, perpetuating the “rare species modelling paradox” (Lomba *et al.* 2010). This could mean that we are ultimately underestimating losses of total species richness because it is precisely these rare species that are most vulnerable to climate change (Ohlemüller *et al.* 2008; Pacifici *et al.* 2015). However, limiting analyses to species with adequate data is a common component of many analyses, including those predicting 30-50% loss in the Neotropics (Wisz *et al.* 2008; Feeley and Silman 2010).

The model predicts at the scale of the data, i.e. the community level, allowing a comparison of a species likelihood of presence in a plot with full uncertainty. However, the model does not explicitly consider mechanistic changes in recruitment, carbon enrichment, seedling survival, or changes in dispersal. As such, the model does not account for plasticity that may allow for species to occur in plots outside its current climate space. Although such mechanistic understanding is needed, the type and resolution of data currently available make this impossible. Finally, this study was limited to the country of Gabon (267,667 km^2^), and may not be comparable to the diverse terrain contained in the 2,250 km2 assessed in the Neotropics (Feeley and Silman 2010).

This study demonstrates that community forecasts are not generalizable across regions, and more studies are needed in understudied biomes like the Afrotropics. Nascent data sets (Enquist *et al.* 2017; Fyllas *et al.* 2017), increased availability of high quality remote sensing (Patterson and Healey 2015; Stavros *et al.* 2017; Silva *et al.* 2018), and new statistical techniques capable of synthesizing multiple types of data (Clark *et al.* 2017) will help in further resolving the responses of the world’s ecosystems. This study serves as an important counterpoint to work done in the Neotropics by providing contrasting predictions for Afrotropical forests with substantially different ecological, evolutionary, and anthropogenic histories. Even though we are predicting a comparatively small reduction in species richness, the effects reported here will have ramifications for whole food webs (Clark, Nuñez and Tomasek, in press; Dirzo *et al.* 2014), and potentially threaten the ecosystem services on which humans depend (McCann 2000; Hooper *et al.* 2005; Balvanera *et al.* 2006; Cardinale *et al.* 2012; Schweiger *et al.* 2018). The differences exposed by this work should serve as motivation for future research using fine scale data to compare the differing responses of tropical biomes to global change.

## References

Abatzoglou JT, Dobrowski SZ, Parks SA, Hegewisch KC. 2018. TerraClimate, a high-resolution global dataset of monthly climate and climatic water balance from 1958-2015. Scientific Data 5.

Asefi-Najafabady S, Saatchi S. 2013. Response of African humid tropical forests to recent rainfall anomalies. Philosophical Transactions of the Royal Society B: Biological Sciences 368.

Balvanera P, Pfisterer AB, Buchmann N, et al. 2006. Quantifying the evidence for biodiversity effects on ecosystem functioning and services. Ecology Letters 9: 1146–1156.

Bongers F, Poorter L, Hawthorne WD, Sheil D. 2009. The intermediate disturbance hypothesis applies to tropical forests, but disturbance contributes little to tree diversity. Ecology Letters 12: 798–805.

Bush MB. 2002. Distributional change and conservation on the Andean flank: A palaeoecological perspective. Global Ecology and Biogeography 11: 463–473.

Bush MB, Silman MR, Urrego DH. 2004. 48,000 Years of Climate and Forest Change in a Biodiversity Hot Spot. Science 303: 827–829.

Cardinale BJ, Duffy JE, Gonzalez A, et al. 2012. Biodiversity loss and its impact on humanity. Nature 486: 59–67.

Carlson BS, Koerner SE, Medjibe VP, White LJT, Poulsen JR. 2017. Deadwood stocks increase with selective logging and large tree frequency in Gabon. Global Change Biology 23: 1648–1660.

Chazdon RL, Finegan B, Capers RS, et al. 2010. Composition and dynamics of functional groups of trees during tropical forest succession in northeastern Costa Rica. Biotropica 42: 31–40.

Clark JS, Nemergut D, Seyednasrollah B, Turner PJ, Zhang S. 2017. Generalized joint attribute modeling for biodiversity analysis: Median-zero, multivariate, multifarious data. Ecological Monographs 87: 34–56.

Colwell RK, Brehm G, Cardelús CL, Gilman AC, Longino JT. 2008. Global warming, elevational range shifts, and lowland biotic attrition in the wet tropics. Science 322: 258–261.

Corlett RT, Primack RB. 2011. Many Tropical Rain Forests In: Tropical Rain Forests.1–31.

Davies SJ, Semui H. 2006. Competitive dominance in a secondary successional rain-forest community in Borneo. Journal of Tropical Ecology 22: 53–64.

Dexter KG, Pennington RT, Oliveira-Filho AT, Bueno ML, Silva de Miranda PL, Neves DM. 2018. Inserting Tropical Dry Forests Into the Discussion on Biome Transitions in the Tropics. Frontiers in Ecology and Evolution 6.

Dirzo R, Young HS, Galetti M, Ceballos G, Isaac NJB, Collen B. 2014. Defaunation in the Anthropocene. Science 345: 401–406.

Engelbrecht BMJ, Comita LS, Condit R, et al. 2007. Drought sensitivity shapes species distribution patterns in tropical forests. Nature 447: 80–82.

Enquist BJ, Bentley LP, Shenkin A, et al. 2017. Assessing trait-based scaling theory in tropical forests spanning a broad temperature gradient. Global Ecology and Biogeography 26: 1357–1373.

Fayolle A, Engelbrecht B, Freycon V, et al. 2012. Geological substrates shape tree species and trait distributions in African moist forests. PLoS ONE 7.

Feeley KJ, Silman MR. 2010. Biotic attrition from tropical forests correcting for truncated temperature niches. Global Change Biology 16: 1830–1836.

Feeley KJ, Silman MR, Bush MB, et al. 2011. Upslope migration of Andean trees. Journal of Biogeography 38: 783–791.

Finegan B. 1996. Pattern and process in neotropical secondary rain forests: The first 100 years of succession. Trends in Ecology and Evolution 11: 119–124.

Fyllas NM, Bentley LP, Shenkin A, et al. 2017. Solar radiation and functional traits explain the decline of forest primary productivity along a tropical elevation gradient. Ecology Letters 20: 730–740.

Gardner TA, Barlow J, Parry LW, Peres CA. 2007. Predicting the uncertain future of tropical forest species in a data vacuum. Biotropica 39: 25–30.

Gond V, Fayolle A, Pennec A, et al. 2013. Vegetation structure and greenness in Central Africa from Modis multi-temporal data. Philosophical Transactions of the Royal Society B: Biological Sciences 368.

Gorelick N, Hancher M, Dixon M, Ilyushchenko S, Thau D, Moore R. 2017. Google Earth Engine: Planetary-scale geospatial analysis for everyone. Remote Sensing of Environment 202: 18–27.

Haffer J. 1969. Speciation in amazonian forest birds. Science 165: 131–137.

Hansen MC, DeFries RS. 2004. Detecting long-term global forest change using continuous fields of tree-cover maps from 8-km Advanced Very High Resolution Radiometer (AVHRR) data for the years 1982-99. Ecosystems 7: 695–716.

Hardwick K, Elliott S. 2016. Second Growth: The Promise of Tropical Rain Forest Regeneration in the Age of Deforestation. Chicago and London: University of Chicago Press.

Hooper DU, Adair EC, Cardinale BJ, et al. 2012. A global synthesis reveals biodiversity loss as a major driver of ecosystem change. Nature 486: 105–108.

Hooper DU, Chapin FS, Ewel JJ, et al. 2005. Effects of biodiversity on ecosystem functioning: A consensus of current knowledge. Ecological Monographs 75: 3–35.

James R, Washington R, Rowell DP. 2013. Implications of global warming for the climate of African rainforests. Philosophical Transactions of the Royal Society B: Biological Sciences 368.

Leal ME. 2009. The past protecting the future: Locating climatically stable forests in West and Central Africa. International Journal of Climate Change Strategies and Management 1: 92–99.

Lomba A, Pellissier L, Randin C, et al. 2010. Overcoming the rare species modelling paradox: A novel hierarchical framework applied to an Iberian endemic plant. Biological Conservation 143: 2647–2657.

Maley J. 1996. The African rain forest - Main characteristics of changes in vegetation and climate from the Upper Cretaceous to the Quaternary. Proceedings of the Royal Society of Edinburgh Section B: Biological Sciences 104: 31–73.

Malhi Y, Adu-Bredu S, Asare RA, Lewis SL, Mayaux P. 2013. African rainforests: Past, present and future. Philosophical Transactions of the Royal Society B: Biological Sciences 368: 20120312–20120312.

Malhi Y, Gardner TA, Goldsmith GR, Silman MR, Zelazowski P. 2014. Tropical Forests in the Anthropocene.

Malhi Y, Wright J. 2004. Spatial patterns and recent trends in the climate of tropical rainforest regions In: Philosophical Transactions of the Royal Society B: Biological Sciences.311–329.

Maslin M, Malhi Y, Phillips O, Cowling S. 2005. New views on an old forest: Assessing the longevity, resilience and future of the Amazon rainforest. Transactions of the Institute of British Geographers 30: 477–499.

Mayaux P, Pekel JF, Desclée B, et al. 2013. State and evolution of the African rainforests between 1990 and 2010. Philosophical Transactions of the Royal Society B: Biological Sciences 368.

McCann KS. 2000. The diversity-stability. Nature 405: 228–233.

Nuñez CL, Clark JS, Clark CJ, Poulsen JR. 2019. Low-intensity logging and hunting have long-term effects on seed dispersal but not fecundity in Afrotropical forests. AoB PLANTS 11.

Ohlemüller R, Anderson BJ, Araújo MB, et al. 2008. The coincidence of climatic and species rarity: High risk to small-range species from climate change. Biology Letters 4: 568–572.

Oslisly R, White L, Bentaleb I, et al. 2013. Climatic and cultural changes in the west Congo Basin forests over the past 5000 years. Philosophical Transactions of the Royal Society B: Biological Sciences 368.

Pacifici M, Foden WB, Visconti P, et al. 2015. Assessing species vulnerability to climate change. Nature Climate Change 5: 215–225.

Pan Y, Birdsey RA, Fang J, et al. 2011. A large and persistent carbon sink in the world’s forests. Science 333: 988–993.

Parmentier I, Malhi Y, Senterre B, et al. 2007. The odd man out? Might climate explain the lower tree α-diversity of African rain forests relative to Amazonian rain forests? Journal of Ecology 95: 1058–1071.

Patterson P, Healey S. 2015. Global ecosystem dynamics investigation (GEDI) LiDAR sampling strategy In: Pushing boundaries: new directions in inventory techniques and applications: Forest Inventory and Analysis (FIA) symposium 2015.245.

Phillips OL, Baker TR. 2002. Field Manual for plot establishment and remeasurement (RAINFOR).

Pimm SL, Raven P. 2000. Extinction by numbers. Nature 403: 843–845.

Poorter L, van der Sande MT, Arets EJMM, et al. 2017. Biodiversity and climate determine the functioning of Neotropical forests. Global Ecology and Biogeography 26: 1423–1434.

Poulsen JR, Koerner SE, Miao Z, Medjibe VP, Banak LN, White LJT. 2017. Forest structure determines the abundance and distribution of large lianas in Gabon. Global Ecology and Biogeography 26: 472–485.

Raich JW, Khoon GW. 1990. Effects of canopy openings on tree seed germination in a malaysian dipterocarp forest. Journal of Tropical Ecology 6: 203–217.

Saatchi S, Asefi-Najafabady S, Malhi Y, et al. 2013. Persistent effects of a severe drought on Amazonian forest canopy. Proceedings of the National Academy of Sciences of the United States of America 110: 565–570.

Sala OE, Chapin FS, Armesto JJ, et al. 2000. Global biodiversity scenarios for the year 2100. Science 287: 1770–1774.

Sannier C, McRoberts RE, Fichet LV, Makaga EMK. 2014. Using the regression estimator with landsat data to estimate proportion forest cover and net proportion deforestation in gabon. Remote Sensing of Environment 151: 138–148.

Schweiger AK, Cavender-Bares J, Townsend PA, et al. 2018. Plant spectral diversity integrates functional and phylogenetic components of biodiversity and predicts ecosystem function. Nature Ecology and Evolution 2: 976–982.

Silva CA, Saatchi S, Garcia M, et al. 2018. Comparison of Small-and Large-Footprint Lidar Characterization of Tropical Forest Aboveground Structure and Biomass: A Case Study from Central Gabon. IEEE Journal of Selected Topics in Applied Earth Observations and Remote Sensing 11: 3512–3526.

Slik JWF, Arroyo-Rodríguez V, Aiba SI, et al. 2015. An estimate of the number of tropical tree species. Proceedings of the National Academy of Sciences of the United States of America 112: 7472–7477.

Stavros EN, Schimel D, Pavlick R, et al. 2017. ISS observations offer insights into plant function. Nature Ecology and Evolution 1.

Sullivan MJP, Talbot J, Lewis SL, et al. 2017. Diversity and carbon storage across the tropical forest biome. Scientific Reports 7: 39102.

van Vuuren DP, Sala OE, Pereira HM. 2006. The future of vascular plant diversity under four global scenarios. Ecology and Society 11.

Wade AM, Richter DD, Medjibe VP, et al. 2019. Estimates and determinants of stocks of deep soil carbon in Gabon, Central Africa. Geoderma 341: 236–248.

Walker K, Sleath E. 2017. A systematic review of the current knowledge regarding revenge pornography and non-consensual sharing of sexually explicit media. Aggression and Violent Behavior 36: 9–24.

Washington R, James R, Pearce H, Pokam WM, Moufouma-Okia W. 2013. Congo basin rainfall climatology: Can we believe the climate models? Philosophical Transactions of the Royal Society B: Biological Sciences 368: 20120296–20120296.

Whitmore TC. 1989. Canopy Gaps and the Two Major Groups of Forest Trees. Ecology 70: 536–538.

Willis KJ, Bennett KD, Burrough SL, Macias-Fauria M, Tovar C. 2013. Determining the response of African biota to climate change: Using the past to model the future. Philosophical Transactions of the Royal Society B: Biological Sciences 368.

Wisz MS, Hijmans RJ, Li J, et al. 2008. Effects of sample size on the performance of species distribution models. Diversity and Distributions 14: 763–773.

