## Appendix A for "Afrotropical Tree Communities May Have Distinct Responses to Forecasted Climate Change"

### A1: Estimated species richness and SE for all plots, 2020-2099.

| | Estimated Plot Species Richness | | | | | | | |
|---|---|---|---|---|---|---|---|---|
| | Wet Model, RCP 4.5 | | Wet Model, RCP 8.5 | | Dry Model, RCP 4.5 | | Dry Model, RCP 8.5 | |
| Year | Mean | SE | Mean | SE | Mean | SE | Mean | SE |
| 2020 | 24.92 | 3.90 | 25.00 | 3.91 | 24.46 | 3.91 | 24.22 | 3.89 |
| 2021 | 24.42 | 3.90 | 24.27 | 3.88 | 24.53 | 3.90 | 24.03 | 3.88 |
| 2022 | 24.46 | 3.90 | 24.73 | 3.87 | 24.17 | 3.89 | 23.76 | 3.86 |
| 2023 | 24.54 | 3.89 | 24.32 | 3.87 | 24.03 | 3.88 | 23.62 | 3.86 |
| 2024 | 24.74 | 3.89 | 23.93 | 3.87 | 24.08 | 3.88 | 23.93 | 3.87 |
| 2025 | 24.50 | 3.89 | 24.09 | 3.86 | 23.94 | 3.87 | 24.04 | 3.87 |
| 2026 | 24.58 | 3.87 | 24.10 | 3.84 | 23.86 | 3.87 | 23.63 | 3.84 |
| 2027 | 24.25 | 3.87 | 23.78 | 3.84 | 23.97 | 3.88 | 23.49 | 3.85 |
| 2028 | 24.79 | 3.87 | 24.23 | 3.84 | 23.51 | 3.85 | 23.00 | 3.81 |
| 2029 | 24.48 | 3.88 | 23.91 | 3.83 | 23.92 | 3.87 | 23.04 | 3.82 |
| 2030 | 24.58 | 3.87 | 23.84 | 3.83 | 23.90 | 3.87 | 23.33 | 3.83 |
| 2031 | 24.59 | 3.86 | 23.62 | 3.83 | 23.50 | 3.85 | 23.13 | 3.82 |
| 2032 | 24.19 | 3.85 | 23.33 | 3.82 | 23.50 | 3.85 | 23.12 | 3.82 |
| 2033 | 23.64 | 3.85 | 23.47 | 3.81 | 23.41 | 3.84 | 22.79 | 3.80 |
| 2034 | 23.93 | 3.86 | 23.55 | 3.80 | 23.30 | 3.84 | 22.59 | 3.79 |
| 2035 | 24.01 | 3.86 | 23.46 | 3.81 | 23.49 | 3.85 | 22.74 | 3.79 |
| 2036 | 23.83 | 3.86 | 23.09 | 3.79 | 23.33 | 3.84 | 22.40 | 3.78 |
| 2037 | 23.79 | 3.82 | 23.59 | 3.80 | 23.10 | 3.83 | 22.92 | 3.80 |
| 2038 | 23.67 | 3.83 | 23.30 | 3.80 | 23.07 | 3.82 | 22.58 | 3.79 |
| 2039 | 23.87 | 3.83 | 22.80 | 3.78 | 23.10 | 3.82 | 22.62 | 3.77 |
| 2040 | 23.69 | 3.83 | 23.18 | 3.77 | 23.21 | 3.83 | 22.26 | 3.76 |
| 2041 | 23.63 | 3.82 | 22.92 | 3.76 | 23.38 | 3.83 | 22.35 | 3.76 |
| 2042 | 23.78 | 3.81 | 22.68 | 3.75 | 23.36 | 3.83 | 21.91 | 3.73 |
| 2043 | 23.74 | 3.80 | 22.58 | 3.73 | 22.86 | 3.80 | 21.80 | 3.72 |
| 2044 | 23.68 | 3.80 | 22.84 | 3.72 | 22.75 | 3.79 | 21.62 | 3.71 |
| 2045 | 23.72 | 3.82 | 22.40 | 3.73 | 22.86 | 3.81 | 21.87 | 3.73 |
| 2046 | 23.41 | 3.83 | 22.83 | 3.72 | 23.30 | 3.83 | 21.90 | 3.73 |
| 2047 | 23.37 | 3.80 | 22.62 | 3.71 | 22.62 | 3.79 | 21.77 | 3.70 |
| 2048 | 23.55 | 3.80 | 22.62 | 3.70 | 22.91 | 3.81 | 21.56 | 3.69 |
| 2049 | 22.96 | 3.79 | 22.31 | 3.70 | 22.42 | 3.77 | 21.40 | 3.69 |
| 2050 | 23.42 | 3.78 | 22.33 | 3.68 | 22.57 | 3.78 | 21.27 | 3.68 |
| 2051 | 23.30 | 3.78 | 22.29 | 3.67 | 22.59 | 3.79 | 21.22 | 3.68 |

| | | | | | | | | |
|---|---|---|---|---|---|---|---|---|
| 2052 | 23.16 | 3.76 | 22.00 | 3.67 | 22.50 | 3.77 | 21.28 | 3.68 |
| 2053 | 22.85 | 3.76 | 21.86 | 3.65 | 22.38 | 3.76 | 21.04 | 3.66 |
| 2054 | 23.75 | 3.78 | 21.77 | 3.64 | 22.39 | 3.77 | 20.86 | 3.64 |
| 2055 | 23.08 | 3.75 | 21.55 | 3.63 | 21.98 | 3.74 | 21.07 | 3.64 |
| 2056 | 23.21 | 3.78 | 22.38 | 3.65 | 22.56 | 3.78 | 21.02 | 3.65 |
| 2057 | 23.55 | 3.80 | 22.40 | 3.63 | 22.81 | 3.80 | 20.81 | 3.62 |
| 2058 | 23.07 | 3.76 | 21.52 | 3.62 | 22.25 | 3.76 | 20.61 | 3.62 |
| 2059 | 22.69 | 3.74 | 21.62 | 3.62 | 22.14 | 3.75 | 20.85 | 3.63 |
| 2060 | 23.14 | 3.74 | 21.56 | 3.61 | 21.76 | 3.73 | 20.69 | 3.62 |
| 2061 | 23.23 | 3.75 | 20.99 | 3.59 | 21.97 | 3.74 | 20.45 | 3.60 |
| 2062 | 22.54 | 3.73 | 21.02 | 3.59 | 21.93 | 3.74 | 20.44 | 3.60 |
| 2063 | 22.32 | 3.73 | 20.78 | 3.56 | 22.07 | 3.74 | 20.39 | 3.58 |
| 2064 | 22.10 | 3.74 | 21.68 | 3.58 | 22.02 | 3.74 | 20.17 | 3.57 |
| 2065 | 22.79 | 3.73 | 21.22 | 3.57 | 21.75 | 3.72 | 20.44 | 3.59 |
| 2066 | 22.69 | 3.73 | 20.70 | 3.54 | 21.88 | 3.73 | 20.07 | 3.55 |
| 2067 | 22.51 | 3.74 | 20.82 | 3.54 | 21.76 | 3.73 | 20.03 | 3.55 |
| 2068 | 22.44 | 3.74 | 20.35 | 3.53 | 21.94 | 3.74 | 20.14 | 3.55 |
| 2069 | 22.70 | 3.73 | 20.32 | 3.51 | 21.83 | 3.73 | 19.65 | 3.52 |
| 2070 | 22.49 | 3.71 | 20.76 | 3.52 | 21.86 | 3.73 | 19.74 | 3.53 |
| 2071 | 22.89 | 3.71 | 20.67 | 3.50 | 21.58 | 3.71 | 19.83 | 3.51 |
| 2072 | 22.86 | 3.72 | 20.51 | 3.49 | 21.61 | 3.71 | 19.58 | 3.50 |
| 2073 | 22.66 | 3.71 | 20.62 | 3.47 | 21.66 | 3.72 | 19.52 | 3.49 |
| 2074 | 22.02 | 3.70 | 20.02 | 3.45 | 21.60 | 3.71 | 19.46 | 3.50 |
| 2075 | 22.13 | 3.71 | 20.26 | 3.47 | 21.65 | 3.71 | 19.57 | 3.48 |
| 2076 | 22.24 | 3.71 | 20.31 | 3.46 | 21.85 | 3.72 | 19.49 | 3.46 |
| 2077 | 22.61 | 3.71 | 20.63 | 3.47 | 21.53 | 3.70 | 19.21 | 3.47 |
| 2078 | 22.22 | 3.70 | 19.87 | 3.43 | 21.37 | 3.69 | 19.15 | 3.45 |
| 2079 | 22.29 | 3.70 | 19.71 | 3.41 | 21.51 | 3.70 | 19.22 | 3.45 |
| 2080 | 22.38 | 3.69 | 19.69 | 3.42 | 21.52 | 3.69 | 19.16 | 3.42 |
| 2081 | 22.22 | 3.68 | 19.77 | 3.41 | 21.23 | 3.68 | 19.17 | 3.42 |
| 2082 | 22.21 | 3.70 | 19.87 | 3.41 | 21.50 | 3.70 | 19.13 | 3.43 |
| 2083 | 22.61 | 3.71 | 19.88 | 3.41 | 21.58 | 3.71 | 19.16 | 3.43 |
| 2084 | 22.98 | 3.71 | 19.71 | 3.39 | 21.48 | 3.70 | 18.83 | 3.40 |
| 2085 | 22.43 | 3.70 | 19.18 | 3.38 | 21.52 | 3.70 | 18.87 | 3.43 |
| 2086 | 22.01 | 3.69 | 19.16 | 3.37 | 21.44 | 3.70 | 18.73 | 3.41 |
| 2087 | 22.12 | 3.69 | 19.36 | 3.35 | 21.68 | 3.71 | 18.64 | 3.38 |
| 2088 | 22.91 | 3.70 | 19.27 | 3.35 | 21.55 | 3.70 | 18.56 | 3.37 |
| 2089 | 22.17 | 3.68 | 19.30 | 3.36 | 21.57 | 3.69 | 18.70 | 3.38 |
| 2090 | 22.16 | 3.69 | 19.61 | 3.35 | 21.30 | 3.69 | 18.52 | 3.37 |
| 2091 | 22.23 | 3.69 | 19.68 | 3.34 | 21.42 | 3.69 | 18.24 | 3.35 |

| | | | | | | | | |
|---|---|---|---|---|---|---|---|---|
| 2092 | 21.72 | 3.68 | 19.66 | 3.34 | 21.20 | 3.68 | 18.55 | 3.34 |
| 2093 | 22.06 | 3.70 | 19.42 | 3.33 | 21.61 | 3.70 | 18.60 | 3.38 |
| 2094 | 22.25 | 3.71 | 19.25 | 3.33 | 21.70 | 3.71 | 18.59 | 3.37 |
| 2095 | 22.80 | 3.71 | 19.48 | 3.33 | 21.58 | 3.71 | 18.39 | 3.36 |
| 2096 | 22.43 | 3.68 | 19.05 | 3.30 | 21.26 | 3.68 | 18.42 | 3.33 |
| 2097 | 22.20 | 3.70 | 19.30 | 3.30 | 21.49 | 3.70 | 18.21 | 3.34 |
| 2098 | 22.12 | 3.69 | 19.31 | 3.30 | 21.41 | 3.69 | 18.52 | 3.35 |
| 2099 | 22.34 | 3.67 | 19.15 | 3.30 | 21.29 | 3.68 | 18.26 | 3.31 |

**A2: Predicted plot species richness in 2020 and 2099, as well as net change and standard error.**

| | Net Change in Species Richness | | | |
|---|---|---|---|---|
| Forest Type | 2020 | 2099 | change | SE |
| | Dry Model, RCP 8.5 | | | |
| Savanna | 25.66 | 18.81 | -6.85 | 3.38 |
| Coastal | 24.49 | 18.24 | -6.25 | 3.32 |
| Aucoumea | 24.43 | 18.24 | -6.19 | 3.31 |
| Congolian | 23.97 | 18.18 | -5.79 | 3.29 |
| | Dry Model, RCP 4.5 | | | |
| Savanna | 25.56 | 22.33 | -3.23 | 3.75 |
| Coastal | 24.33 | 21.18 | -3.16 | 3.67 |
| Aucoumea | 24.46 | 21.20 | -3.26 | 3.67 |
| Congolian | 24.09 | 21.15 | -2.94 | 3.66 |
| | Wet Model, RCP 8.5 | | | |
| Savanna | 26.24 | 19.69 | -6.54 | 3.38 |
| Coastal | 25.07 | 19.14 | -5.93 | 3.32 |
| Aucoumea | 25.01 | 19.10 | -5.91 | 3.29 |
| Congolian | 24.76 | 18.98 | -5.78 | 3.27 |
| | Wet Model, RCP 4.5 | | | |
| Savanna | 26.25 | 23.91 | -2.34 | 3.77 |
| Coastal | 24.87 | 22.42 | -2.45 | 3.70 |
| Aucoumea | 25.11 | 22.52 | -2.58 | 3.68 |
| Congolian | 24.67 | 22.07 | -2.59 | 3.64 |

### A3: The models used in the climate prediction

| Precipitation | Maximium Temperature | Minimum Temperature | |
|---|---|---|---|
| NorESM1.M | CSIRO.Mk3.6.0 | CSIRO.Mk3.6.0 | **Highest Five Models** |
| MRI.ESM.MR | ACCESS1.0 | ACCESS1.0 | |
| MPI.ESM.LR | CanESM2 | CanESM2 | |
| MIROC5 | IPSL.CM5A.LR | IPSL.CM5A.LR | |
| IPSL.CM5A.MR | IPSL.CM5A.MR | IPSL.CM5A.MR | |
| MIROC.ESM | MIROC.ESM | MIROC.ESM | **Lowest Five Models** |
| MIROC.ESM.CHEM | MIROC.ESM.CHEM | MIROC.ESM.CHEM | |
| inmcm4 | inmcm4 | inmcm4 | |
| bcc.csm1.1 | MRI.CGCM3 | MRI.CGCM3 | |
| CanESM2 | CNRM.CM5 | NorESM1.M | |

**A4: Predicted change in species abundance per plot by end of century for both wet and dry models, as well as RCP 4.5, and RCP 8.5.**

| | Estimated Species Abundance | | | | | | | |
|---|---|---|---|---|---|---|---|---|
| | Wet Model, RCP 4.5 | | Wet Model, RCP 8.5 | | Dry Model, RCP 4.5 | | Dry Model, RCP 8.5 | |
| Species Name | Mean | SE | Mean | SE | Mean | SE | Mean | SE |
| *Diospyros spp.* | 5.38 | 1.41 | 21.82 | 2.09 | 8.18 | 1.92 | 24.45 | 1.92 |
| *Aucoumea klaineana* | 16.05 | 4.95 | 25.97 | 10.22 | 3.56 | 4.01 | 11.67 | 2.32 |
| *Staudtia gabonensis* | 4.3 | 1.03 | 10.32 | 2.15 | 2.59 | 0.89 | 9.44 | 0.85 |
| *Desbordesia glaucescens* | 1.67 | 0.64 | 5.57 | 1.13 | 1.33 | 0.56 | 5.31 | 0.8 |
| *Coula edulis* | 1.69 | 1.01 | 4.45 | 1.26 | 1.15 | 0.93 | 4.21 | 0.88 |
| *Strombosia spp.* | 1.65 | 0.62 | 4.55 | 1.41 | 0.7 | 0.51 | 3.79 | 0.93 |
| *Pausinystalia johimbe* | 0.98 | 0.53 | 2.74 | 0.77 | 0.69 | 0.45 | 2.43 | 0.6 |
| *Macaranga spp.* | -0.19 | 0.49 | 0.66 | 0.89 | 0.68 | 0.85 | 1.48 | 0.75 |
| *Mytragyna ciliata* | 0.87 | 0.52 | 2.34 | 0.67 | 0.56 | 0.59 | 1.41 | 0.6 |
| *Dialium pachyphyllum* | -0.57 | 0.53 | 0.22 | 0.71 | 0.53 | 0.79 | 0.69 | 0.71 |
| *Scytopetalum klaineanum* | 1.37 | 0.57 | 3.3 | 0.97 | 0.48 | 0.55 | 2.26 | 0.53 |
| *Xylopia aethiopica* | 2.31 | 0.94 | 4.06 | 1.69 | 0.42 | 0.66 | 1.97 | 0.64 |
| *Irvingia gabonensis* | 0.15 | 0.55 | 1.09 | 0.65 | 0.4 | 0.6 | 1.1 | 0.67 |
| *Nauclea diderrichii* | -0.12 | 0.37 | 1.1 | 0.66 | 0.28 | 0.58 | 1.07 | 0.53 |
| *Diogoa zenkeri* | 0.82 | 0.7 | 1.37 | 0.93 | 0.23 | 0.79 | 0.84 | 0.71 |
| *Polyalthia spp.* | -0.36 | 0.56 | -0.37 | 0.67 | 0.18 | 0.73 | 0.18 | 0.72 |
| *Trichoscypha acuminata* | -0.3 | 0.39 | 0.04 | 0.59 | 0.09 | 0.49 | 0.34 | 0.53 |
| *Zanthoxylum heitzii* | 0.52 | 0.44 | 1.4 | 0.61 | 0.09 | 0.34 | 0.67 | 0.38 |
| *Maprounea membranacea* | 0.65 | 0.59 | 0.9 | 0.75 | -0.05 | 0.43 | 0.26 | 0.4 |
| *Ongokea gore* | 0 | 0.42 | 0.48 | 0.48 | -0.06 | 0.39 | 0.26 | 0.45 |
| *Annickia chlorantha* | 0.73 | 0.51 | 0.8 | 0.65 | -0.08 | 0.36 | 0.05 | 0.31 |
| *Odyendyea gabonensis* | 0.11 | 0.38 | 0.12 | 0.55 | -0.08 | 0.41 | 0.04 | 0.42 |
| *Grewia coriacea* | 0.17 | 0.42 | 0.54 | 0.5 | -0.1 | 0.39 | 0.4 | 0.45 |
| *Baphia spp.* | -1.32 | 0.64 | -1.67 | 0.82 | -0.11 | 0.88 | -1.8 | 0.76 |
| *Xylopia spp.* | 0.13 | 0.31 | 0.1 | 0.37 | -0.17 | 0.3 | 0.07 | 0.24 |
| *Klainedoxa gabonensis* | 0.36 | 0.39 | -0.09 | 0.62 | -0.2 | 0.32 | -0.23 | 0.38 |

| | | | | | | | | |
|---|---|---|---|---|---|---|---|---|
| *Klainedoxa spp.* | -0.47 | 0.4 | -0.54 | 0.46 | -0.26 | 0.53 | -0.57 | 0.49 |
| *Barteria fistulosa* | -0.15 | 0.43 | -0.71 | 0.57 | -0.3 | 0.48 | -1.06 | 0.49 |
| *Pycnanthus angolensis* | 1.38 | 0.83 | 0.97 | 1.34 | -0.3 | 0.72 | -0.24 | 0.55 |
| *Dacryodes buettneri* | -0.31 | 0.58 | -0.75 | 0.66 | -0.32 | 0.58 | -0.49 | 0.66 |
| *Dacryodes normandii* | -0.19 | 0.32 | -0.42 | 0.37 | -0.36 | 0.31 | -0.63 | 0.32 |
| *Dacryodes igaganga* | -0.52 | 0.36 | -0.79 | 0.45 | -0.39 | 0.38 | -0.84 | 0.56 |
| *Scyphocephalium ochocoa* | 0.57 | 0.83 | -0.1 | 1.1 | -0.41 | 0.68 | -0.45 | 0.58 |
| *Dialium spp.* | -0.34 | 0.46 | -1.1 | 0.49 | -0.42 | 0.5 | -1.02 | 0.55 |
| *Distemonanthus benthamianus* | -0.67 | 0.38 | -1.05 | 0.52 | -0.44 | 0.4 | -1.02 | 0.53 |
| *Anonidium mannii* | -0.43 | 0.41 | -0.69 | 0.64 | -0.47 | 0.43 | -1.12 | 0.98 |
| *Trichilia spp.* | -0.73 | 0.49 | -0.85 | 0.68 | -0.5 | 0.7 | -0.78 | 0.64 |
| *Centroplacus glaucinus* | -0.58 | 0.4 | -0.86 | 0.49 | -0.5 | 0.49 | -0.71 | 0.51 |
| *Anthonotha spp.* | -0.59 | 0.37 | -0.95 | 0.46 | -0.51 | 0.45 | -0.77 | 0.53 |
| *Erismadelphus exsul* | -0.32 | 0.37 | -1.06 | 0.49 | -0.51 | 0.31 | -0.85 | 0.42 |
| *Chrysophyllum spp.* | -0.52 | 0.3 | -1.12 | 0.47 | -0.55 | 0.32 | -0.99 | 0.43 |
| *Piptadeniastrum africanum* | -0.21 | 0.38 | -0.9 | 0.48 | -0.56 | 0.4 | -0.82 | 0.38 |
| *Pterocarpus soyauxii* | -0.61 | 0.44 | -1.28 | 0.47 | -0.59 | 0.43 | -1.14 | 0.52 |
| *Pausinystalia macroceras* | -1.31 | 0.58 | -1.85 | 0.98 | -0.63 | 0.74 | -1.9 | 0.86 |
| *Afrostyrax lepidophyllus* | -0.77 | 0.34 | -1.03 | 0.48 | -0.65 | 0.49 | -1.62 | 0.73 |
| *Pseudospondias macrocarpa* | -0.68 | 0.4 | -1.34 | 0.52 | -0.7 | 0.4 | -1.36 | 0.54 |
| *Polyalthia suaveolens* | -0.17 | 0.5 | -1.07 | 0.63 | -0.72 | 0.5 | -1.14 | 0.46 |
| *Panda oleosa* | -0.97 | 0.35 | -1.44 | 0.61 | -0.73 | 0.55 | -2 | 0.83 |
| *Tetraberlinia bifoliolata* | -0.51 | 0.46 | -1.44 | 0.81 | -0.74 | 0.42 | -1.06 | 0.64 |
| *Xylopia staudtii* | -0.69 | 0.39 | -1.42 | 0.48 | -0.76 | 0.38 | -1.31 | 0.53 |
| *Symphonia globulifera* | -0.74 | 0.47 | -1.42 | 0.86 | -0.79 | 0.49 | -0.97 | 0.66 |
| *Beilschmiedia spp.* | -0.94 | 0.42 | -1.85 | 0.52 | -0.82 | 0.55 | -1.56 | 0.6 |
| *Dacryodes macrophylla* | -0.92 | 0.61 | -1.52 | 0.88 | -0.84 | 0.57 | -1.25 | 0.75 |
| *Erythrophleum ivorense* | -1.03 | 0.55 | -1.52 | 0.58 | -0.84 | 0.44 | -1.54 | 0.58 |
| *Duboscia macrocarpa* | -0.72 | 0.34 | -1.41 | 0.55 | -0.84 | 0.36 | -1.46 | 0.55 |
| *Strombosia pustulata* | -0.69 | 0.56 | -1.73 | 0.68 | -0.86 | 0.48 | -1.64 | 0.67 |
| *Carapa procera* | -0.79 | 0.49 | -1.87 | 0.58 | -0.89 | 0.52 | -1.69 | 0.64 |
| *Pentaclethra macrophylla* | -1.14 | 0.49 | -1.97 | 0.82 | -0.92 | 0.48 | -2.09 | 0.8 |

| | | | | | | | | |
|---|---|---|---|---|---|---|---|---|
| *Pentaclethra eetveldeana* | -1.86 | 0.71 | -3.01 | 1.26 | -0.92 | 0.76 | -3.16 | 0.97 |
| *Drypetes spp.* | -0.62 | 0.47 | -1.85 | 0.66 | -0.99 | 0.45 | -1.69 | 0.65 |
| *Klaineanthus gaboniae* | -1.3 | 0.58 | -2.5 | 0.74 | -1.14 | 0.55 | -2.52 | 0.89 |
| *Garcinia spp.* | -0.81 | 0.82 | -2.57 | 1.55 | -1.16 | 0.72 | -1.59 | 1.01 |
| *Trichoscypha spp.* | -1.1 | 0.6 | -2.42 | 0.88 | -1.21 | 0.53 | -2.14 | 0.82 |
| *Celtis tessmannii* | -1.32 | 0.57 | -2.12 | 0.89 | -1.22 | 0.53 | -2.79 | 1.19 |
| *Uapaca spp.* | -1.01 | 0.68 | -2.33 | 1.02 | -1.29 | 0.71 | -1.82 | 0.87 |
| *Scorodolphloeus zenkeri* | -1.33 | 0.6 | -2.08 | 1.16 | -1.3 | 0.88 | -3.62 | 1.52 |
| *Strombosiopsis tetrandra* | -0.67 | 0.75 | -2.42 | 0.79 | -1.33 | 0.65 | -2.63 | 0.75 |
| *Hymenostegia pellegrinii* | -1.13 | 0.67 | -1.62 | 0.9 | -1.35 | 0.77 | -2.61 | 1.3 |
| *Dacryodes spp.* | -1.25 | 0.75 | -2.67 | 1.19 | -1.35 | 0.74 | -2.16 | 1.06 |
| *Cola spp.* | -1.15 | 0.71 | -3.11 | 0.85 | -1.44 | 0.62 | -2.96 | 0.82 |
| *Petersianthus macrocarpus* | -1.37 | 0.92 | -2.03 | 1.39 | -1.58 | 0.99 | -2.85 | 1.92 |
| *Heisteria parvifolia* | -1.56 | 0.7 | -3.76 | 1.57 | -1.87 | 0.89 | -2.96 | 1.3 |
| *Coelocaryon preussii* | -3.41 | 1.67 | -6.64 | 3.61 | -3.43 | 1.71 | -4.66 | 3.05 |
| *Dichostemma glaucescens* | -2.24 | 3.17 | -6.85 | 3.31 | -4.23 | 3.23 | -7.39 | 2.92 |
| *Plagiostyles africana* | -6.18 | 3.16 | -9.16 | 5.48 | -6.21 | 2.7 | -11.64 | 6.28 |
| *Santiria trimera* | -7.17 | 2.15 | -14.69 | 4.9 | -7.52 | 2.08 | -14.67 | 4.78 |

**A5: Posterior parameter chains show convergence.**

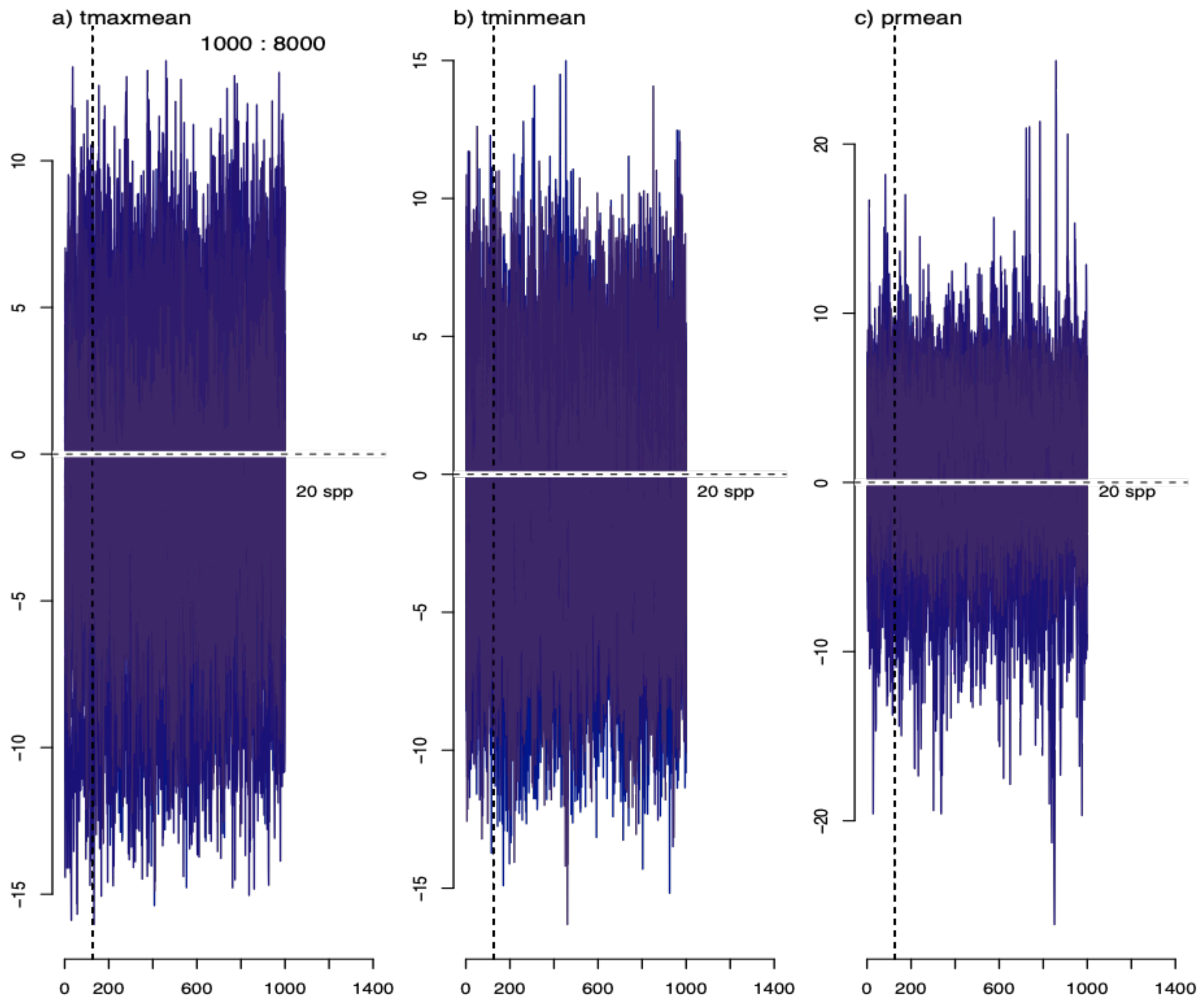

**A6: A comparison of long-term average climate space for all 104 plots compared to predicted climate space in 2099 from all models (wet and dry) and both scenarios (RCP 4.5 and 8.5).**

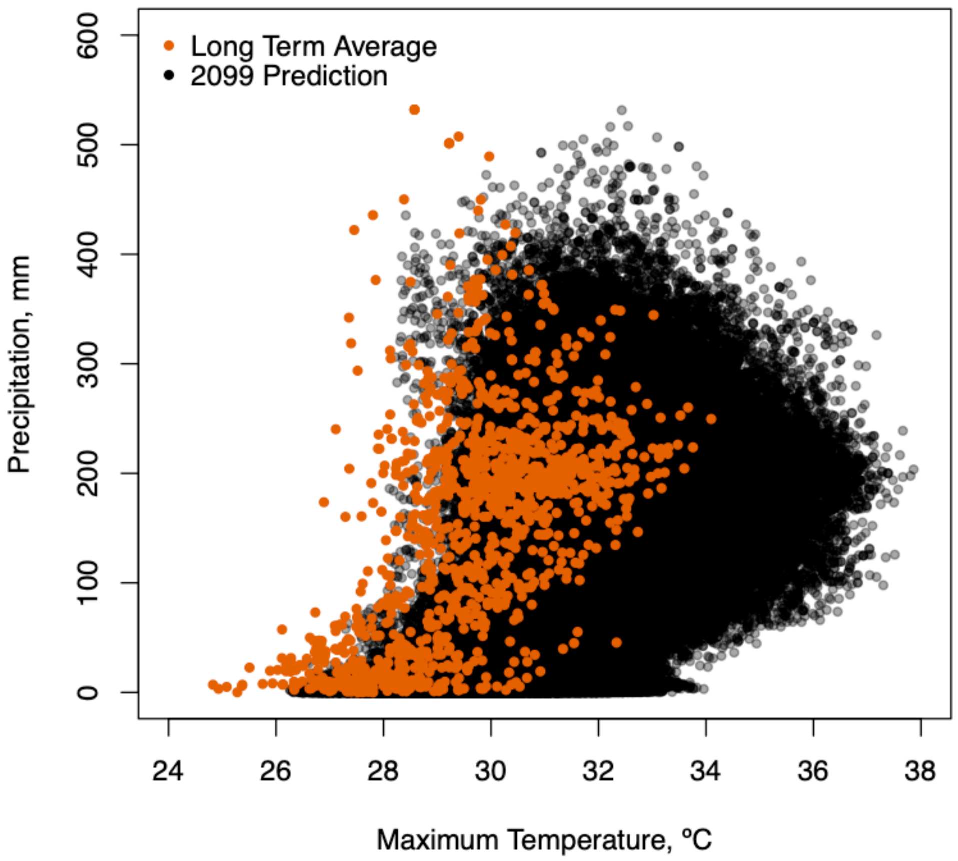

**A7: A comparison of predicted vs. observed values discrete species abundance confirms model fit.**

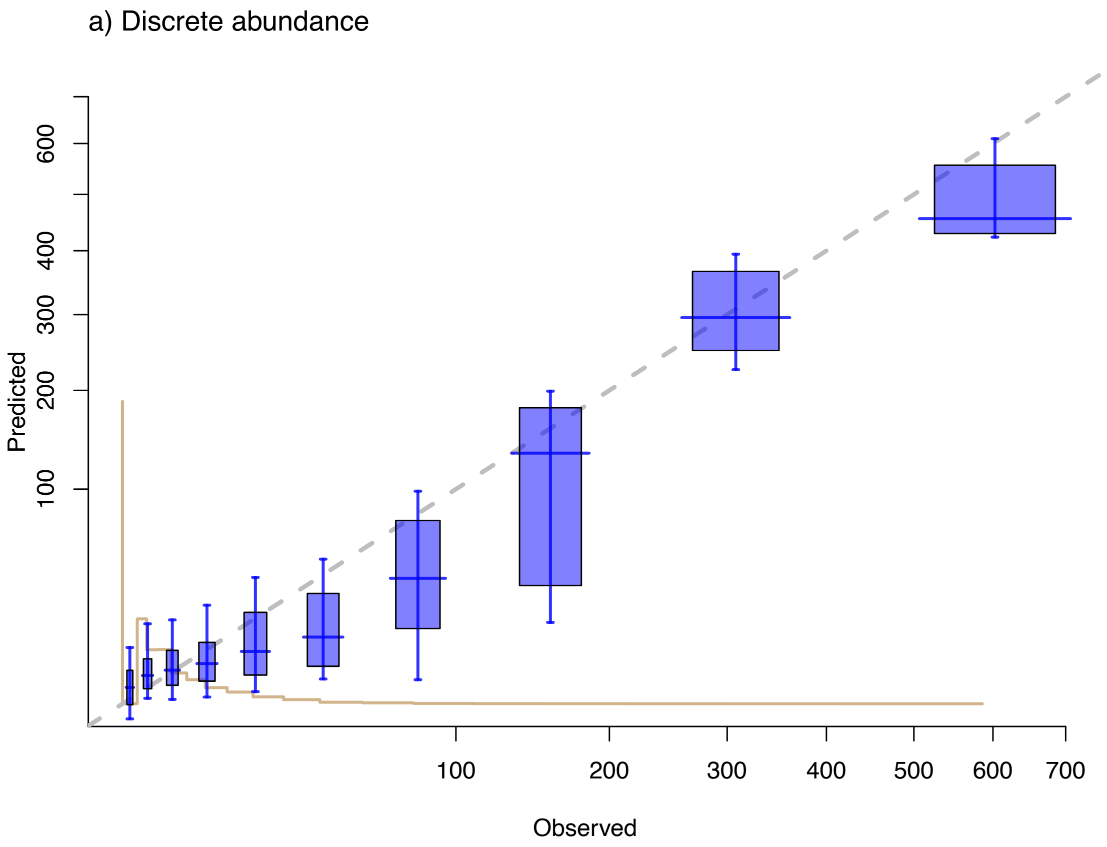
